# SE(3) Equivalent Graph Attention Network as an Energy-Based Model for Protein Side Chain Conformation

**DOI:** 10.1101/2022.09.05.506704

**Authors:** Deqin Liu, Sheng Chen, Shuangjia Zheng, Sen Zhang, Yuedong Yang

## Abstract

Protein design energy functions have been developed over decades by leveraging physical forces approximation and knowledge-derived features. However, manual feature engineering and parameter tuning might suffer from knowledge bias. Learning potential energy functions fully from crystal structure data is promising to automatically discover unknown or high-order features that contribute to the protein’s energy. Here we proposed a graph attention network as an energy-based model for protein conformation, namely GraphEBM. GraphEBM is equivariant to the SE(3) group transformation, which is the important principle of modern machine learning for molecules-related tasks. GraphEBM was benchmarked on the rotamer recovery task and outperformed both Rosetta and the state-of-the-art deep learning based methods. Furthermore, GraphEBM also yielded promising results on combinatorial side chain optimization, improving 22.2% *χ*_1_ rotamer recovery to the PULCHRA method on average.

## I. Introduction

Proteins are chain-like polymers composed of a sequence of dehydration condensed amino acids. Most native proteins folded into stable conformations. According to Anfinsen’s thermodynamic hypothesis, the native state is the one with the lowest free energy [1], [2]. This hypothesis inspired the application of potential energy functions in protein structure prediction [3]–[5] and protein rational design [6]–[9]. The direct optimization of physical energy function composed of complex force fields suffered from the rough energy landscape [10]. Therefore, researchers have developed the statistical potential methods [4], [11], [12] that data-driven fit energetic terms combined with physically motivated force fields. After several decades of development, to-date energy functions for protein design have incorporated extensive feature engineering, extracting physical and biochemical knowledge-based features that contributed to the protein’s energy [13], [14]. Deep learning has been shown to have the ability to capture the hidden high-order dependencies between source and target [15]. A number of deep-learning based methods including our previous works have successfully leveraged the deep learning methods in the field of protein design [16]–[18] protein engineering [19], [20] and protein structure prediction [21], [22]. Therefore, it is promising to learn protein energy function fully from crystal structure data by deep learning methods.

Du et.al. took an initial step toward fully learning protein energy function from data [23]. They leveraged Transformer [24] as a energy-based model [25] for protein side chain conformation. Since the major degrees of freedom in protein conformation are the dihedral rotations [26], they evaluated their method on the side chain rotamer recovery task, where a number of side chain conformations are sampled conditioned on the local structure context and the one with lowest predicted energy is picked as the predicted conformation. However, we argue that their architecture is not equivariant to the SE(3) group transformation, which means that their architecture is not guaranteed to output the same energy value after rigid rotation or translation on a protein conformation. SE(3) group equivariance has been a principle of modern machine learning on molecule-related tasks [27], [28]. Several SE(3) equivariant architectures have been developed for protein design [29], [30] but they focused on residue-wise backbone structures instead atomic conformation. The directional message passing neural network (DimeNet) [31], [32] is an atomic resolution SE(3) equivariant architectures for small molecular graphs. However, DimeNet has not been refined for protein-related tasks and it suffered from training gradient exploding for the sampled conformation without physical constraints.

Here we propose GraphEBM, to our best knowledge, the first SE(3) equivariant energy-based model for protein side chain conformation. We tested GraphEBM on the side chain rotamer recovery task through two different sampling strategies. On average, for both sampling strategies, GraphEBM outperformed two well-known energy function of Rosetta [14], [33] and the state-of-the-art deep learning based method [23]. As a further study, we then applied GraphEBM on combinatorial side chain optimization for a fixed backbone [34], [35]. Starting from the protein conformation yielded by PULCHRA [36], we simply adopted a naive strategy to solve the combinatorial optimization problem. To this end, 22.2% optimized side chain conformations were obtained by GraphEBM on average, showing the potential application of GraphEBM on the general problem settings of protein rational design.

To summarize, our contributions are as follows:

- We proposed the first SE(3) equivariant energy based model for protein side chain conformation.
- We refined the massage aggregating architecture of DimeNet by combining it with Graph Attention Network (GAT).
- To overcome the training gradient exploding problem of DimeNet, we added a smooth factor in the Bessel function with theoretically and experimentally analysis of its influence on the performance of gradient descent optimization.

## II. Methods

The goal is to score the side-chain conformation for a given fixed target backbone structure. This section describes the procedure of side-chain scoring, the architecture of the model, the setting of the smooth factor and the training strategy.

### A. Preliminary

Proteins are large biomolecules and macromolecules composed by one or more long chains of amino acid residues. The protein structure can be viewed as a molecule graph with atoms as nodes and covalent bonds as edges. We abstract the conformation as a graph *G* = (*V, E*). *G* is an undirected graph with a set *V* of nodes and a set *E* of edges. The model is to score the graph *G*. Fig 1 shows the graph input and the architecture of GraphEBM. The red color atoms are variable atoms and the orange are the atoms selected, and they with their bonds compose the input graph.

**Fig. 1.**
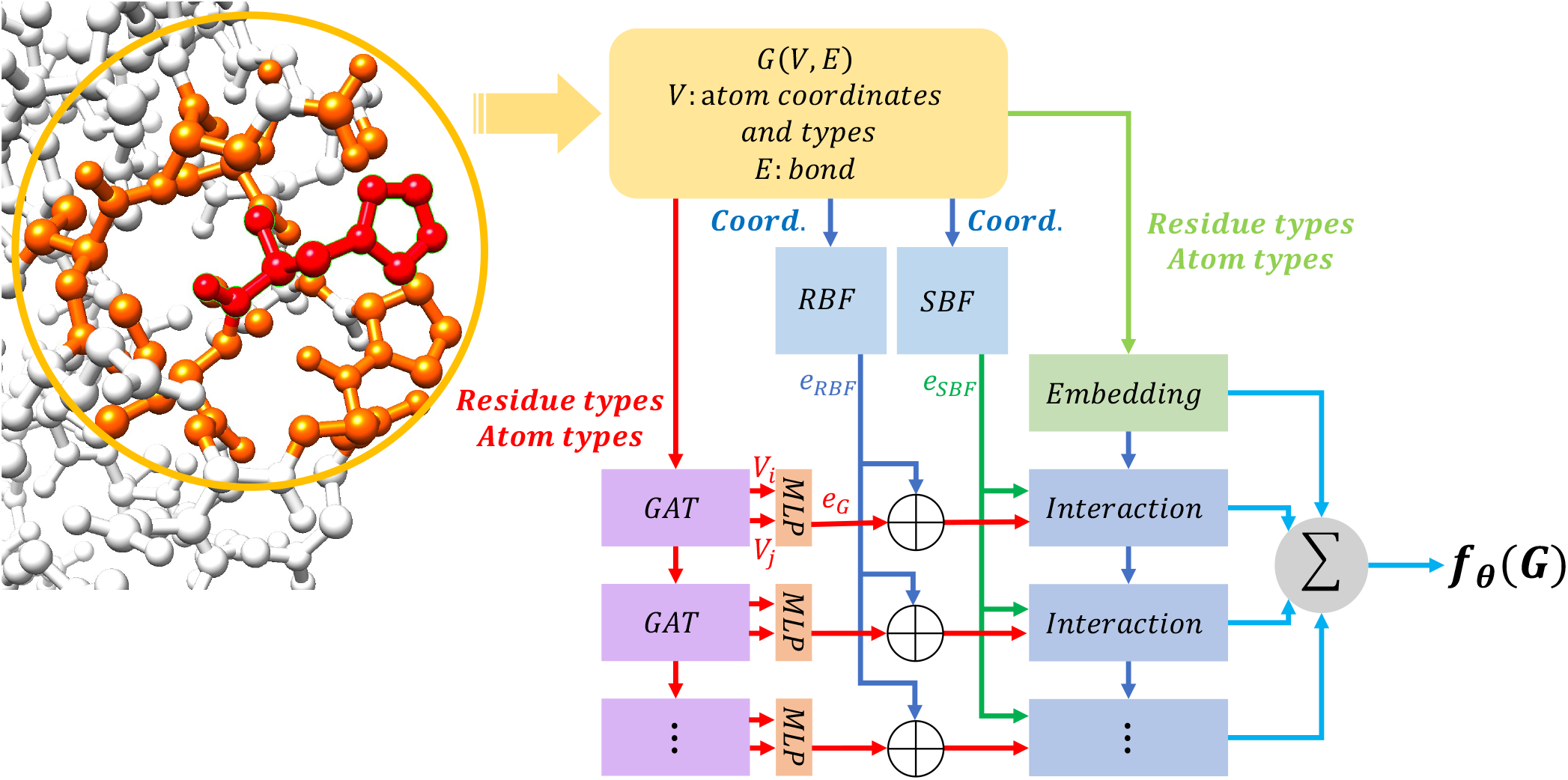
Details on GraphEBM’s architecture: The structural context(≤ 5*Å*) of a given residue is represented by a graph *G* that contains nodes *V* represented by atom types and coordinates and edges *E* represented by *e*_*RBF*_, *e*_*SBF*_ and *e*_*G*_ where *e*_*RBF*_ and *e*_*SBF*_ are calculated by the Bessel functions, *e*_*G*_ is the pair-wise representation of the GAT’s output, and ⊕ denotes the concatenate operation. GraphEBM aggregates the embedding and interaction modules and outputs the final score of the protein conformation.

#### 1) Selection of input nodes

The input graph contains atoms with distance < 5 *Å* to any atom of the conformation of the given residue.

#### 2) Representations of input

The input graph of the local conformation consists of node features: (i) atom types (N, C, O, S); (ii) residue types (which of the 20 types of amino acids the atom belongs to); (iii) atom indicator (indicate if an atom is variable), and atom bonds as edges.

### B. Model architecture

Our model is based on the DimeNet++, which uses the Schrödinger equation and density functional theory. Its features are extracted from the geometry relations by radial basis function(RBF) and the spherical Bessel functions(SBF). RBF using distance and SBF using distance and angles are both SE(3)-Equivariance which avoiding expensive data augmentation strategies [37]. The embedding layer generates the inital message embeddings using the atom types. Then, the interaction module, combination of complex linear layers and directional message passing, update the the embedding from embedding layer or upper network. The interaction module also outputs scalars for the energy of this layer.

Our model GraphEBM described by Fig 1, improves the DimeNet++ and extend its capabilities to the energy prediction of proteins. We update the embedding layer for residue types which can’t be embedded before and introduce GAT and MLP to learn the atom bonds ignored in DimeNet++. The RBF and SBF focus on the distance and angles between atoms. The RBF is defined by

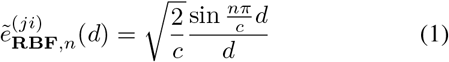

where *n* ∈ [1 … *N*_*RBF*_] denotes the order of RBF, *c* denotes cutoff distance to consider their interactions and *d* denotes the distance between atom *i* and *j*. And the SBF defined by

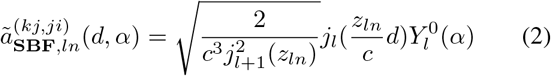

where *l* ∈ [1 … *N*_*SBF*_] denotes the order of Bessel functions, *j*_*l*_ denotes the *l*-order Bessel function, *z*_*ln*_ denotes the *n*-th root of the *l*-order Beseel function and 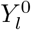denotes the Spherical harmonics. Considering the bonds have an important influence on the physical properties of atoms, we should use a Graph Neural Network to catch the topological structure and residue type information. The attention mechanism has proven to be very effective so far, so we introduce GAT to aggregate the residue types and atom types information by message passing on atom bonds. The GAT can be described by

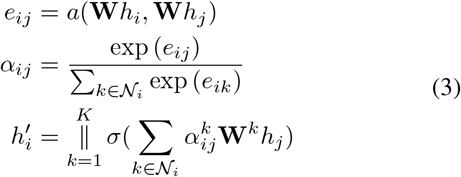

where *h*_*i*_ is the input feature of node *i, a* is a shared attentional mechanism, **W** is a weight matrix, *K* is the number of attention heads, *N*_*i*_ denotes the set of neighbors of node *i, σ* denotes the activate function and ∥ denotes the concatenation operation. We using 8 layers of the interaction module and GAT determined by experiment. When the number of layers is not too high, GAT can continuously update the node representation as the number of layers increases relatively. So with the massage passing of GAT, the node representation can catch bigger field structure information. The output of GAT is ℝ^*N* ×*K*×*H*^, where *N* is the number of nodes, *K* is the number of attention heads and *H* is the number of hidden dimension. We mean the *K* dimension and pair nodes if their edge in *e*_**RBF**_ to 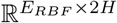 where *E*_*RBF*_ is the number of RBF edges. We use a MultiLayer Perceptron to map the pair-wised embedding to 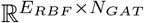 and concatenate with *e*_**RBF**_ to 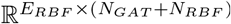. Finally, the energy is the summary of every interaction module output.

### C. Smooth factor

In Fourier-based calculations, the multiplicative inverse of the polynomial is crucial and indispensable for precision. in this work, sampling a side-chain conformation is random, and is not constrained by physical laws. This sampling strategy will generate some atoms which are so close that the Bessel function overflows or causes exploding gradient. Inspired by Laplace smooth, the distance in RBF and SBF can add a *λ* factor for smoothness and stability in training procedures.

The smooth factor can take the model out of this trouble by stabilizing the gradient. The new RBF and SBF is defined as followed. We will discuss the smooth factor selection and analysis later.

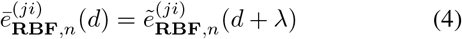

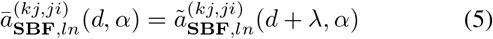

### D. Training and loss function

The model is to score the conformations. But in training procedure, we only have the native conformations which is the state with the lowest free energy according to Anfinsen’s thermodynamic hypothesis. So we sample some negative conformations different from the native state. The loss function can cover this knowledge by using the partition function in statistical mechanics. The partition function can represent the whole system states, so the loss function can just maximize the native conformation’s probability described by the Boltzmann distribution: *p*_*θ*_ = *exp*(−*E*_*θ*_ (*x, c*))*/Z*(*c*), where *Z* = ∫ *exp*(−*E*_*θ*_ (*x, c*))*dx*, where *θ* denotes the learnable parameters, *c* denotes the atoms of the surrounding molecular context and *x* denotes the side chain conformation. In this distribution, partition function means the energy of one molecular formula’s all conformations which can be approximated to conformations generated from the sampling strategy. So, the more negative samples can make the partition function more approach the real system. Furthermore by assuming the *q*(*x*|*c*) distribution is uniform, the loss function can be simplified as followed

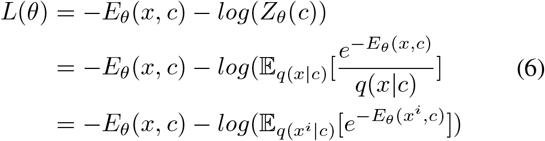

where *q*(*x*|*c*) denotes the conformation probability. After the simplification, the partition function can be approximated by the logarithmic sum of energy of all conformations

## III. Experiments

### A. Dataset

**TABLE I.** Test data summary

| dataset | All | Buried | Medium | Surface |
| --- | --- | --- | --- | --- |
| Test dataset | 10720 | 1263 | 5587 | 3870 |

The unit is the number of amino acids

The dataset is the same with [23], which contains high-resolution PDB structures and removes similar proteins in the test dataset. The training dataset has 12473 structures, and the test dataset has 121 structures.

### B. Evaluation and comprison setting

The energy function should distinguish the conformation closest to the native conformation from samples. The comparison methods consist of Deep Learning methods and the Rosetta energy functions. We rerun the Rosetta to predict the side-chain conformations of the test dataset using Rosetta socre12 and Rosetta ref2015 [14] energy functions. And we compare it with the Atom Transformer [23] which is the state-of-the-art-model in this task.

The Rosetta is the powerful software in protein design, so for comparable with it, the two sampling methods, corresponding to the Rosetta protocol rotamer trials and rotamer trials min, are discrete and continuous sampling strategies. We use the same test sampling strategy as Atom Transformer to reimplement the sampling strategy in Rosetta. Sampling the *µ*(mean) and *σ*(standard deviation) of *χ* angles from the rotamer library needs the backbone *ϕ* and *ψ* angles of the residue. But the rotamer library is a discrete database for backbone angles every 10 degrees and has a weighted combination of *µ* and *σ* of *χ* angles for every 10 degrees backbone angles. Every residue backbone angle can be put in a grid surrounded by the closest discrete point from the rotamer library. Then, samples can be weighted and generated from the grid points by distance or uniform. The discrete strategy is the *χ*_1_ and *χ*_2_ mean and (−1, 0, 1)·*σ* combinations. Another continuous strategy is based on the *µ* and *σ* which can describe a Gaussian distribution 𝒩 (*µ*, 4 · *σ*). This is also the training sampling strategy, but with the uniform sampling for the matched combinations from the rotamer library. The energy function scores every conformation sampled and selects a conformation with the lowest energy. When all *χ* angles of the selected conformation are within 20° of the ground truth, the rotamer is recovered correctly.

For a more detailed analysis, we used the classification from [23] to define buried residues(≥24), medium residues(*others*) and surface residues(<16) by the number of neighbors within 10Å of the residue’s *C*_*β*_.

### C. Result of side chain rotamer recovery

Table II shows the comparison of our model with two versions of Rosetta energy functions and Atom Transformer(reported by [23]). We run Rosetta on the same test dataset of 121 proteins in the rotamer-trials mover and rt-min mover. For a fair comparison, the same or comparable sampling strategies above-mentioned are used to evaluate the model. In table II, our model’s sampling strategy is discrete while rotamer-trials mover and continuous while rt-min mover. Moreover, our model performs better than Rosetta energy functions and the Atom Transformer on both strategies. We split the residues by above-defined types of buried, medium and surface. Our model shows the significant outperformance in surface residue. The surface residue rotamer recovery are more difficult because of the less of physical constraints compared to buried residues. But GraphEBM performs 10% better than other models. We infer this improvement should is due to that GraphEBM additionally introduces the geometry information compared to other methods. Rosetta energy functions are based on the interaction graph [14] which has been calculated and stored as a library. This interaction graph and the same interaction strategy in Atom Transformer focus on the pair-wise relation, but GraphEBM considers the angle and distance between three atoms.

**TABLE II.** Rotamer recovery accuracy over the test dataset

| Discrete sampling strategy |  |  |  |  |
| --- | --- | --- | --- | --- |
| Model | Avg | Buried | Medium | Surface |
| Rosetta score12 (rotamer-trials) | 73.1 | 90.7 | 78.4 | 59.7 |
| Rosetta ref2015 (rotamer-trials) | 75.1 | **91.5** | **80.4** | 62.5 |
| Atom Transformer | 70.2 | 91.3 | 73.7 | 58.2 |
| Atom Transformer (ensemble) | 71.5 | 91.2 | 75.3 | 59.5 |
| GraphEBM | **76.0** | 87.0 | 78.3 | **69.2** |

| Continuous sampling strategy |  |  |  |  |
| --- | --- | --- | --- | --- |
| Model | Avg | Buried | Medium | Surface |
| Rosetta score12 (rt-min) | 73.2 | 91.0 | 78.2 | 60.2 |
| Rosetta ref2015 (rt-min) | 75.8 | **91.9** | 80.8 | 62.5 |
| Atom Transformer | 73.1 | 91.1 | 79.3 | 58.3 |
| Atom Transformer (ensemble) | 74.1 | 91.1 | 80.3 | 59.5 |
| GraphEBM | **78.4** | 90.6 | **81.4** | **70.2** |

Fig 2 reports the recovery rate for every type of residue without the ALA and GLY, because they don’t have any *χ* angles. We run the Rosetta rt-min mover for the test dataset in the default setting without the energy function. The table shows our model mostly outperforms or comes close to other models even the Rosetta ref2015 energy function. Consistent with other methods, the performance of our method on ARG(with a positive charge and polarity), GLN(with polarity), GLU(with a negative charge and polarity), and LYS(with a positive charge and polarity) rotamer recovery is worse than other residues. Furthermore, their hydropathy index ranks in the top four [38]. And those residues are more possible on the surface.

**Fig. 2.**
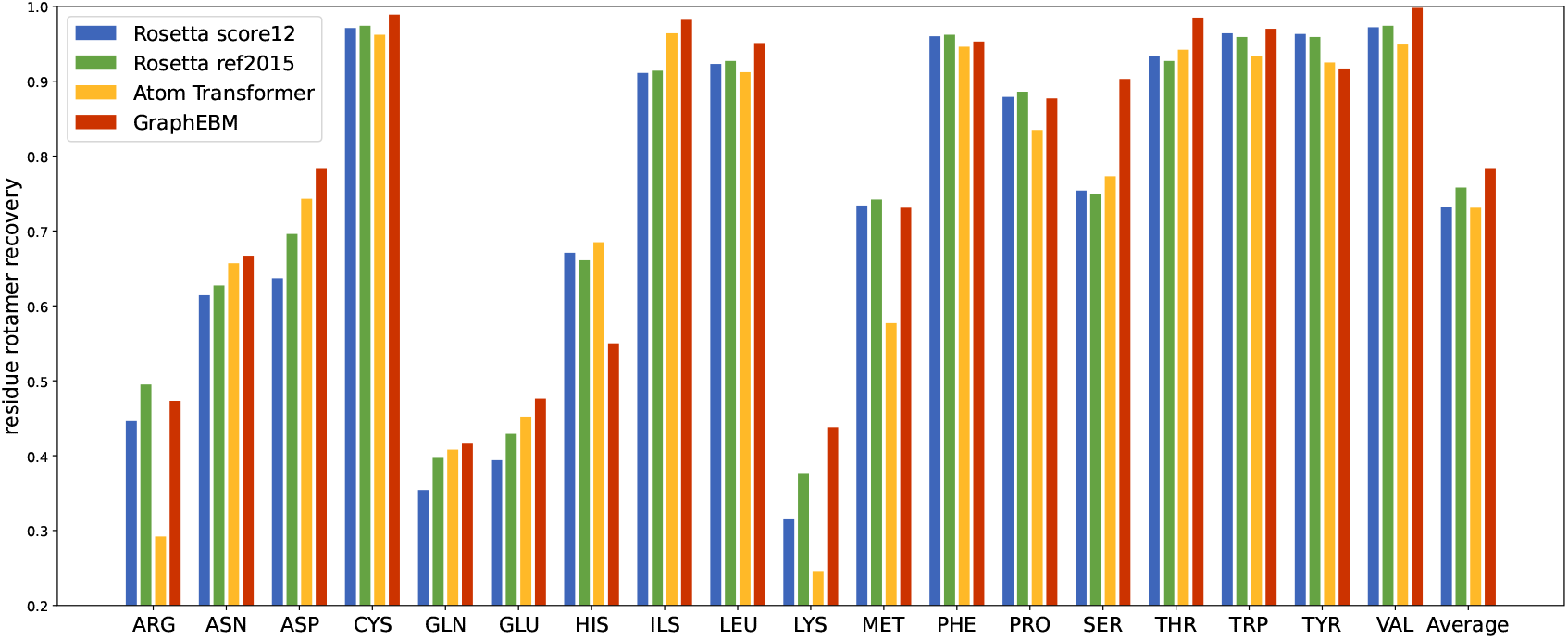
Residues rotamer recovery

### D. The Δχ_1_ and Δχ_2_ angles distribution

Fig 3.a shows the distribution of the Δ*χ*_1_. The *χ*_1_ angle is most precise angle of the native conformation, so we visualize the angle proportion. The performance of GraphEBM is almost close to Rosetta when Δ*χ*_1_ is small. But the Rosetta has more extreme distribution. It is worth noting that the proportion of GraphEBM approaches 1 faster than Rosetta when Δ*χ*_1_ *>* 10°. Fig 3.b has the same trend of Δ*χ*_1_ + Δ*χ*_2_. And those figures show the different sampling strategies have a greater impact on the model accuracy, because the same strategy’s models have the same performance while Δ*χ* is small.

**Fig. 3.**
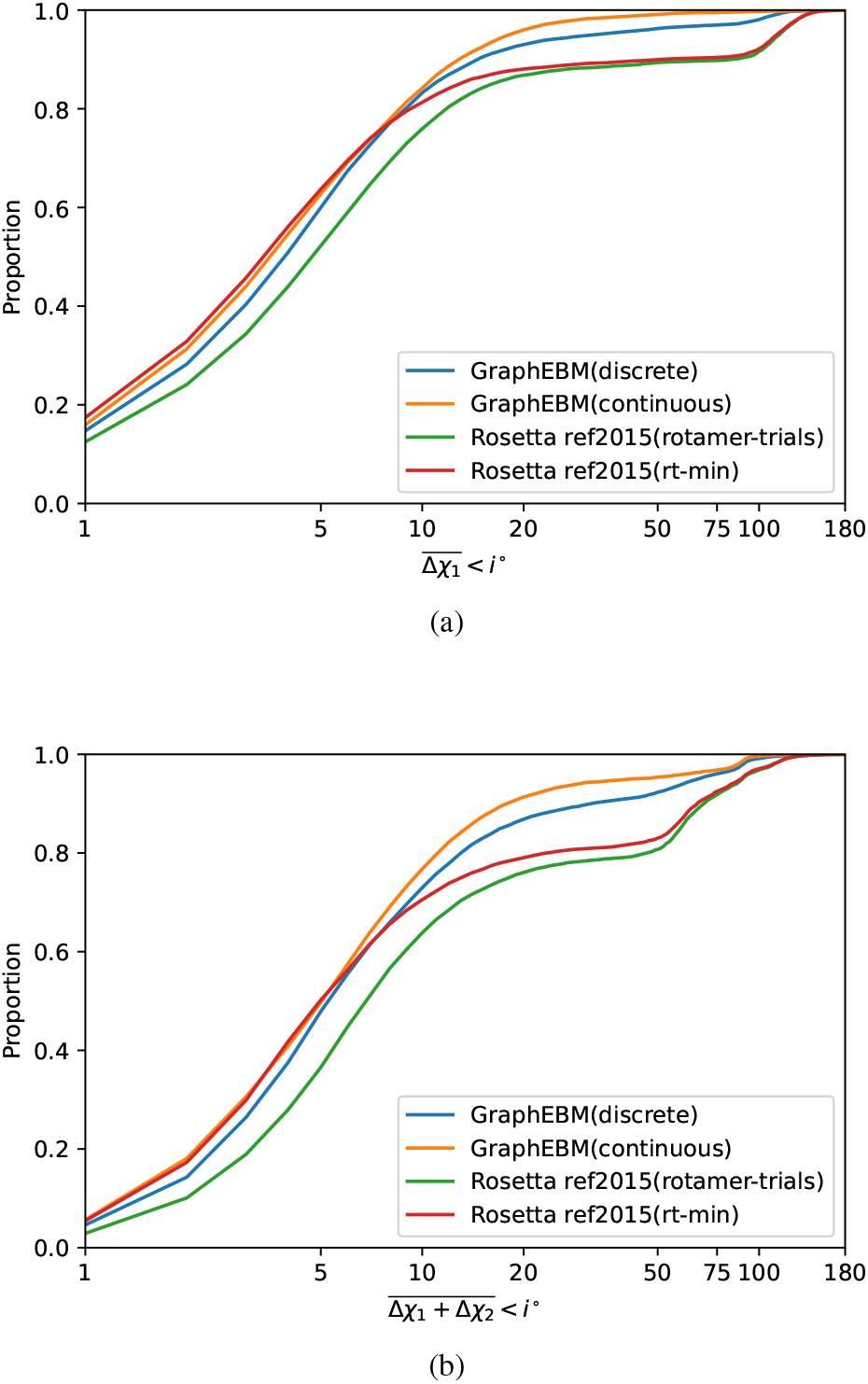
The distribution of Δ*χ*_1_ and Δ*χ*_2_.

## IV. Analysis

### A. Ablation study on smooth factor and graph attention nerual network

For training stability, we introduce a smooth factor in the RBF and SBF. We need to prove the smooth factor can reduce the gradient and analyze its sensitivity. In the RBF(1) and SBF(2), the Bessel function case the gradient explosion.

Equation as followed shows the RBF is the 0-th order Bessel function, so we can focus on the Bessel function’s gradient.

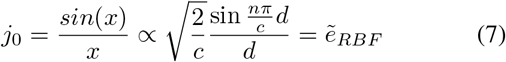

In the gradient-based optimizer [39], the parameter update process is

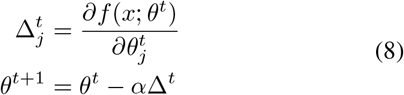

where *θ* denotes parameters of the model *f*, Δ denotes the gradient of the *t*-th iteration and *α* denotes the learning rate. If the input *x* become the Bessel function result, the gradient will become

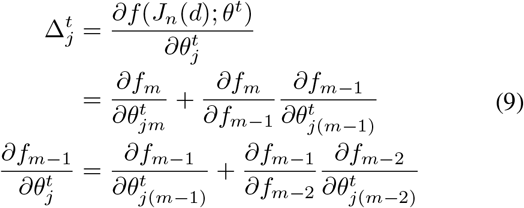

where the first item on the right side denotes the *∂* only operates the *m*-th layer parameters. So we can keep simplifying the problem until the first layer of the model. Without loss of generality, we can specify the first layer as a linear layer. The first layer gradient is calculated by

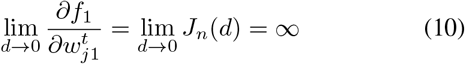

where *w* is the weight of the linear layer, holds when the function *J*_*n*_(*d*) is the Bessel function of the second kind. So if we introduce the smooth factor in the form of Eq (4) and Eq (5), the gradient will reduce. But the introduction of the smooth factor will also reduce the range of the function which cause the resolution of this input descend. Θ denotes the resolution of the Bessel function, Δ denotes the gradient while updating the parameters. The bigger the Θ the model better while the smaller the Δ the model more stable. They can be defined by

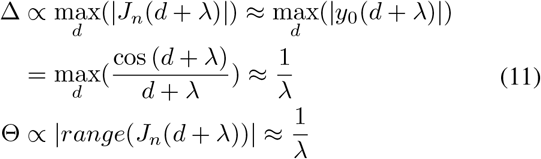

where *range*() denotes the function range. So the performance of the model conditioned by *λ* can be described by the

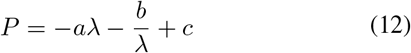

where *P* denotes the performance and *a, b, c* are constants.

Fig 4 shows the same tendency described by Equation (12) with *a* = 0.091, *b* = 0.013, *c* = 0.850 fitted by MATLAB. It demonstrates the proof of the smooth factor’s sensitivity and effect is correct and its results are what we expected.

**Fig. 4.**
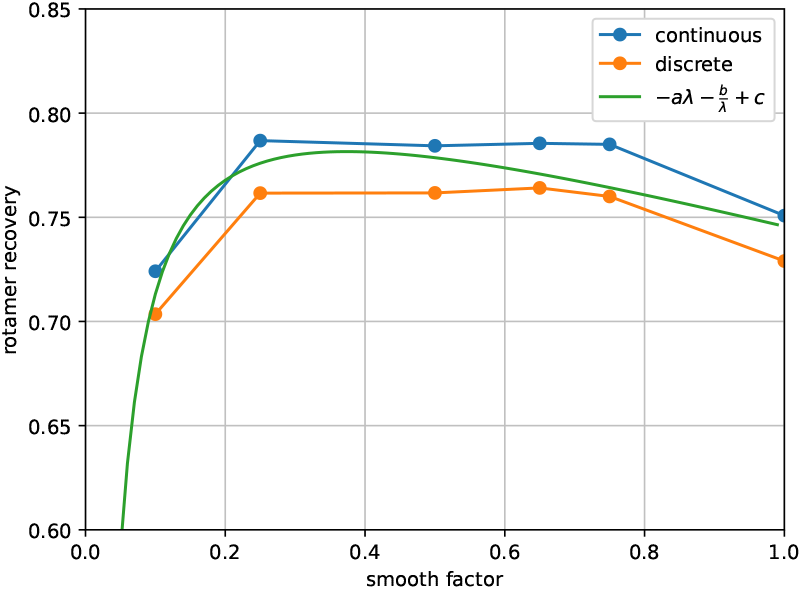
The rotamer recovery rate with the different smooth factor.

We also ran the GraphEBM without smooth factor and graph attention network. We seted a gradient maximum cutoff of 100 to avoid exploding gradient, and got 0.029 relative lower recovery of 0.755 in continuous strategy. Consistently, the ablation of GAT got 0.018 relative lower recovery of 0.766 in the same strategy.

### B. Energy visualization

The buried side chains are more tightly packed [26] than others because of the fewer degrees of freedom. Buried/Medium/Surface energy curve in Fig 5.a shows the steeper response to variations away from the native conformation. Because some residues like Tyr, Asp, and Phe are symmetric about *χ*_2_. Fig 5.b shows a 180° periodicity as the symmetry of them.

**Fig. 5.**
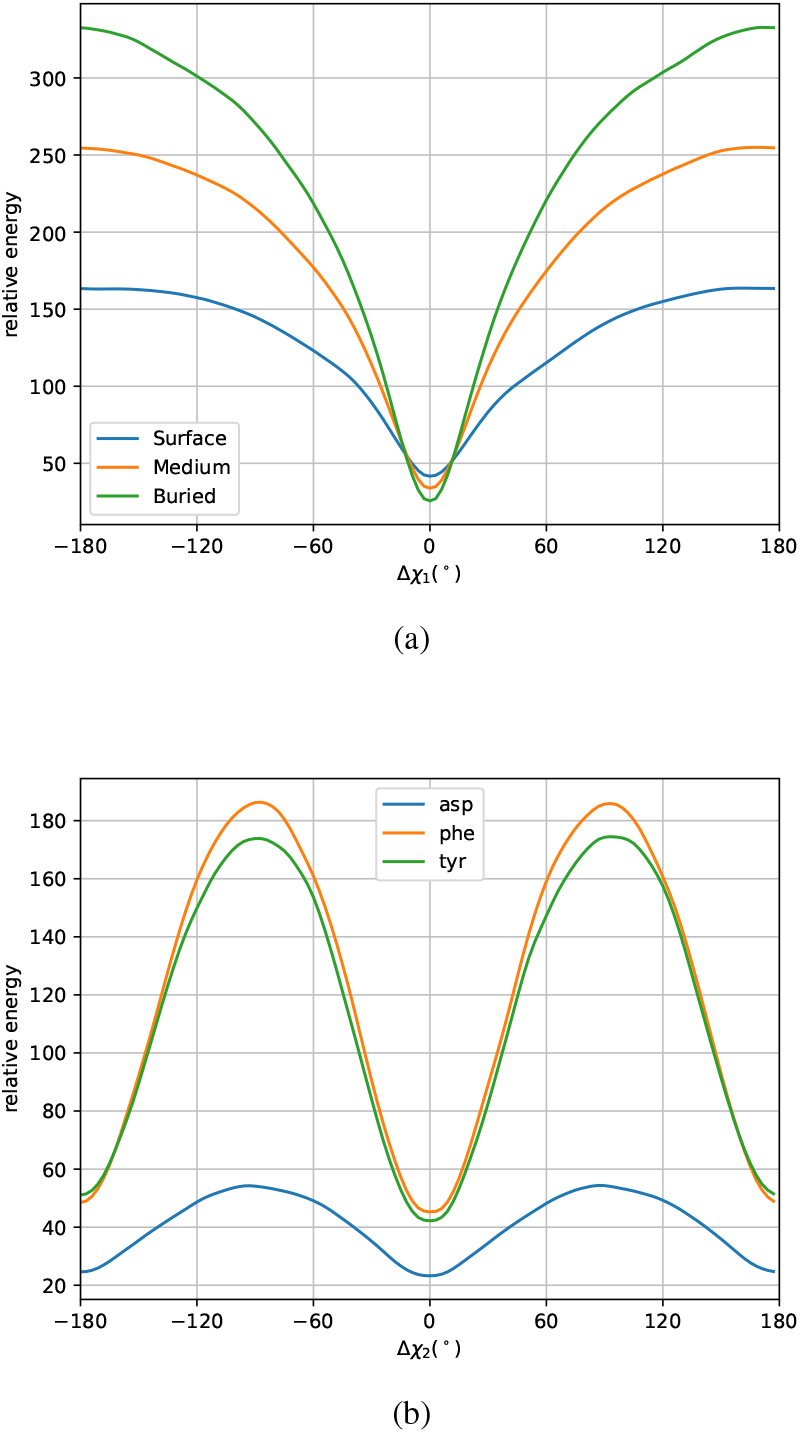
Relative energy curve: a) Different position energy curve; b) Energy curve for the amino acids Asp, Phe, and Tyr with terminal symmetry about *χ*_2_

### C. Combinational side chain optimization

For testing our model on Combinational side chain optimization, we run the PULCHRA [36] to generate an init side-chain conformation with only backbone. PULCHRA is a geometry method and very fast for side chain generation. But, it’s recovery rate are bad even only considering *chi*_1_.The recover strategy is to iterate residue by residue and select the optimal until every residue is stable. GraphEBM trying to recover the side chain from the init conformation by PULCHRA and GraphEBM improve the recovery rate from 53.5% to 75.7% in *χ*_1_ and for 38.3% to 62.2% in *χ*_1_ + *χ*_2_ the same. For our simple iteration strategy, the improvement is sufficiently significant and considerable.

For a more intuitive display, we visualize the surface of a protein(PDBID:1TUKA). In Fig 6, the side-chain from PULCHRA is disorganized and unaligned to the ground truth, but the refined side-chain is more regular and closed to the truth.

**Fig. 6.**
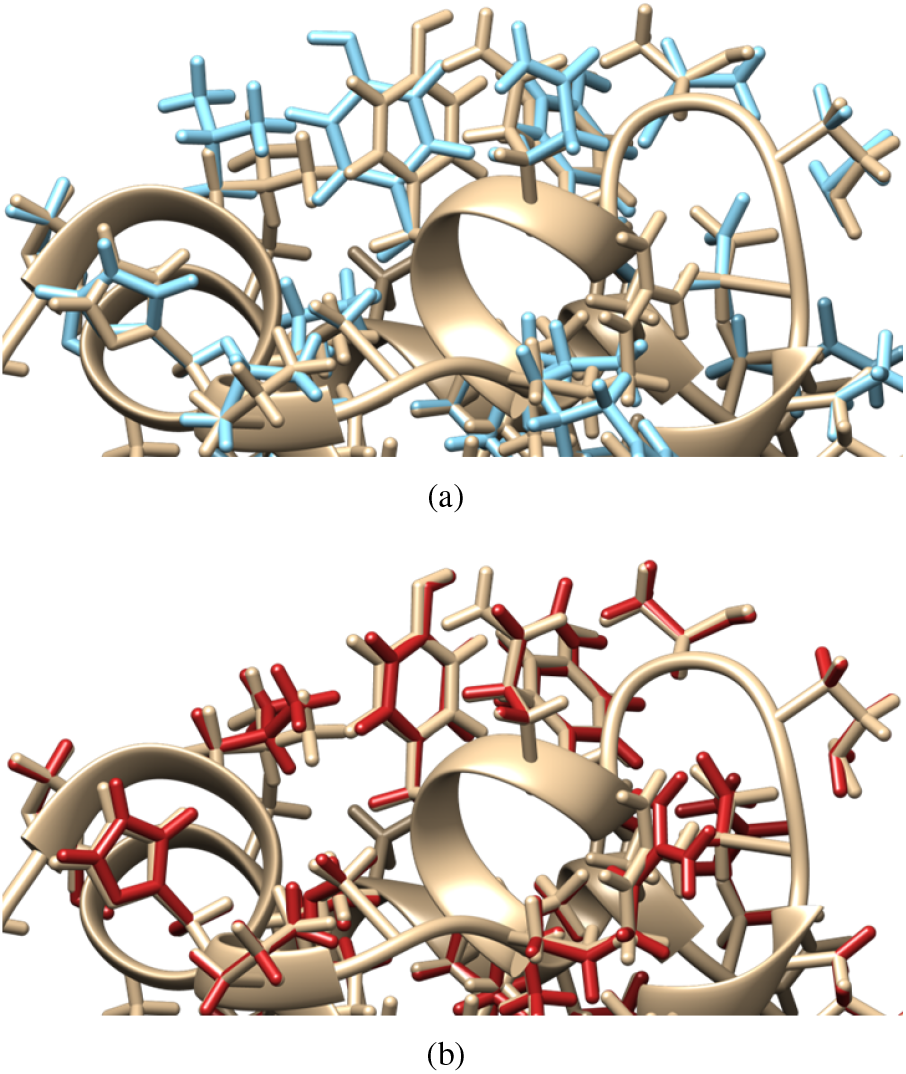
The PDB deposited structure, the PULCHRA predicted all-atom model and the our side chain conformation refined model is colored tan, sky-blue and brick-red respectively.

## V. Conclusion

We propose GraphEBM combining the DimeNet++ and GAT in order to obtain more detailed information from residue types and atom bonds. In the energy-based model training strategy, we introduce a smooth factor for stabilization. And we perform better than Rosetta energy functions in the rotamer recovery task. Those energy functions are based on physical calculation and knowledge. This energy-based strategy can use the simplest knowledge to achieve this performance. GraphEBM plays an essential role in the model because of DFT. We infer the DFT still has a more profound application in Deep Learning. The model trained on recovering one side chain can recover the whole protein’s side chain. But this is limited by the sampling strategy because the whole protein recovery task is a combinatorial optimization problem. Based on simple sampling strategy can not solve that problem well. We think the future work is to generate the side chain conformation.

## VI. Code availability

All code is available at github.com/biomed-AI/GraphEBM.

## VII. Acknowledgment

This study has been supported by the National Key R&D Program of China (2020YFB0204803), National Natural Science Foundation of China (61772566), Guangdong Key Field R&D Plan (2019B020228001 and 2018B010109006), Introducing Innovative and Entrepreneurial Teams (2016ZT06D211), Guangzhou S&T Research Plan (202007030010).

## References

[1] C. B. Anfinsen, “Principles that govern the folding of protein chains,” Science, vol. 181, no. 4096, pp. 223–230, 1973.

[2] C. Anfinsen and H. Scheraga, “Experimental and theoretical aspects of protein folding,” Advances in protein chemistry, vol. 29, pp. 205–300, 1975.

[3] A. Liwo, J. Lee, D. R. Ripoll, J. Pillardy, and H. A. Scheraga, “Protein structure prediction by global optimization of a potential energy function,” Proceedings of the National Academy of Sciences, vol. 96, no. 10, pp. 5482–5485, 1999.

[4] T. Lazaridis and M. Karplus, “Effective energy functions for protein structure prediction,” Current opinion in structural biology, vol. 10, no. 2, pp. 139–145, 2000.

[5] C. A. Rohl, C. E. Strauss, K. M. Misura, and D. Baker, “Protein structure prediction using rosetta,” in Methods in enzymology. Elsevier, 2004, vol. 383, pp. 66–93.

[6] D. B. Gordon, S. A. Marshall, and S. L. Mayo, “Energy functions for protein design.” Current opinion in structural biology, vol. 9, no. 4, pp. 509–513, 1999.

[7] S. M. Lippow and B. Tidor, “Progress in computational protein design,” Current opinion in biotechnology, vol. 18, no. 4, pp. 305–311, 2007.

[8] P.-S. Huang, S. E. Boyken, and D. Baker, “The coming of age of de novo protein design,” Nature, vol. 537, no. 7620, pp. 320–327, 2016.

[9] B. Kuhlman and P. Bradley, “Advances in protein structure prediction and design,” Nature Reviews Molecular Cell Biology, vol. 20, no. 11, pp. 681–697, 2019.

[10] T. Lazaridis and M. Karplus, ““ new view” of protein folding reconciled with the old through multiple unfolding simulations,” Science, vol. 278, no. 5345, pp. 1928–1931, 1997.

[11] S. Tanaka and H. A. Scheraga, “Medium-and long-range interaction parameters between amino acids for predicting three-dimensional structures of proteins,” Macromolecules, vol. 9, no. 6, pp. 945–950, 1976.

[12] M. J. Sippl, “Calculation of conformational ensembles from potentials of mena force: an approach to the knowledge-based prediction of local structures in globular proteins,” Journal of molecular biology, vol. 213, no. 4, pp. 859–883, 1990.

[13] F. E. Boas and P. B. Harbury, “Potential energy functions for protein design,” Current opinion in structural biology, vol. 17, no. 2, pp. 199–204, 2007.

[14] R. F. Alford, A. Leaver-Fay, J. R. Jeliazkov, M. J. O’Meara, F. P. DiMaio, H. Park, M. V. Shapovalov, P. D. Renfrew, V. K. Mulligan, K. Kappel, J. W. Labonte, M. S. Pacella, R. Bonneau, P. Bradley, R. L. Dunbrack, R. Das, D. Baker, B. Kuhlman, T. Kortemme, and J. J. Gray, “The Rosetta All-Atom Energy Function for Macromolecular Modeling and Design,” Journal of Chemical Theory and Computation, vol. 13, no. 6, pp. 3031–3048, 2017.

[15] Y. LeCun, Y. Bengio, and G. Hinton, “Deep learning,” nature, vol. 521, no. 7553, pp. 436–444, 2015.

[16] Z. Li, Y. Yang, E. Faraggi, J. Zhan, and Y. Zhou, “Direct prediction of profiles of sequences compatible with a protein structure by neural networks with fragment-based local and energy-based nonlocal profiles,” Proteins: Structure, Function, and Bioinformatics, vol. 82, no. 10, pp. 2565–2573, 2014.

[17] S. Chen, Z. Sun, L. Lin, Z. Liu, X. Liu, Y. Chong, Y. Lu, H. Zhao, and Y. Yang, “To improve protein sequence profile prediction through image captioning on pairwise residue distance map,” Journal of chemical information and modeling, vol. 60, no. 1, pp. 391–399, 2019.

[18] J. Ingraham, V. Garg, R. Barzilay, and T. Jaakkola, “Generative models for graph-based protein design,” Advances in neural information processing systems, vol. 32, 2019.

[19] E. C. Alley, G. Khimulya, S. Biswas, M. AlQuraishi, and G. M. Church, “Unified rational protein engineering with sequence-based deep representation learning,” Nature methods, vol. 16, no. 12, pp. 1315–1322, 2019.

[20] X. Lv, J. Chen, Y. Lu, Z. Chen, N. Xiao, and Y. Yang, “Accurately predicting mutation-caused stability changes from protein sequences using extreme gradient boosting,” Journal of chemical information and modeling, vol. 60, no. 4, pp. 2388–2395, 2020.

[21] J. Jumper, R. Evans, A. Pritzel, T. Green, M. Figurnov, O. Ronneberger, K. Tunyasuvunakool, R. Bates, A. Ž ídek, A. Potapenko et al., “Highly accurate protein structure prediction with alphafold,” Nature, vol. 596, no. 7873, pp. 583–589, 2021.

[22] Y. Cai, X. Li, Z. Sun, Y. Lu, H. Zhao, J. Hanson, K. Paliwal, T. Litfin, Y. Zhou, and Y. Yang, “Spot-fold: Fragment-free protein structure prediction guided by predicted backbone structure and contact map,” Journal of Computational Chemistry, vol. 41, no. 8, pp. 745–750, 2020.

[23] Y. Du, J. Meier, J. Ma, R. Fergus, and A. Rives, “Energy-based models for atomic-resolution protein conformations,” 2004.13167 [cs, qbio, stat], 2020.

[24] A. Vaswani, N. Shazeer, N. Parmar, J. Uszkoreit, L. Jones, A. N. Gomez, Ł. Kaiser, and I. Polosukhin, “Attention is all you need,” Advances in neural information processing systems, vol. 30, 2017.

[25] Y. LeCun, S. Chopra, R. Hadsell, M. Ranzato, and F. J. Huang, “A Tutorial on Energy-Based Learning,” p. 59.

[26] J. S. Richardson and D. C. Richardson, “Principles and patterns of protein conformation,” in Prediction of protein structure and the principles of protein conformation. Springer, 1989, pp. 1–98.

[27] N. Thomas, T. Smidt, S. Kearnes, L. Yang, L. Li, K. Kohlhoff, and P. Riley, “Tensor field networks: Rotation-and translation-equivariant neural networks for 3d point clouds,” arXiv preprint 1802.08219, 2018.

[28] O.-E. Ganea, X. Huang, C. Bunne, Y. Bian, R. Barzilay, T. S. Jaakkola, and A. Krause, “Independent SE(3)-Equivariant Models for End-to-End Rigid Protein Docking,” in International Conference on Learning Representations, 2021.

[29] B. Jing, S. Eismann, P. Suriana, R. J. Townshend, and R. Dror, “Learning from protein structure with geometric vector perceptrons,” arXiv preprint 2009.01411, 2020.

[30] M. McPartlon, B. Lai, and J. Xu, “A deep se (3)-equivariant model for learning inverse protein folding,” bioRxiv, 2022.

[31] J. Klicpera, J. Groß, and S. Günnemann, “Directional Message Passing for Molecular Graphs,” 2003.03123 [physics, stat], 2020.

[32] J. Klicpera, S. Giri, J. T. Margraf, and S. Günnemann, “Fast and uncertainty-aware directional message passing for non-equilibrium molecules,” arXiv preprint 2011.14115, 2020.

[33] A. Leaver-Fay, M. J. O’Meara, M. Tyka, R. Jacak, Y. Song, E. H. Kellogg, J. Thompson, I. W. Davis, R. A. Pache, S. Lyskov, J. J. Gray, T. Kortemme, J. S. Richardson, J. J. Havranek, J. Snoeyink, D. Baker, and B. Kuhlman, “Scientific Benchmarks for Guiding Macromolecular Energy Function Improvement,” in Methods in Enzymology. Elsevier, 2013, vol. 523, pp. 109–143.

[34] P. Tuffery, C. Etchebest, S. Hazout, and R. Lavery, “A new approach to the rapid determination of protein side chain conformations,” Journal of Biomolecular structure and dynamics, vol. 8, no. 6, pp. 1267–1289, 1991.

[35] L. Holm and C. Sander, “Fast and simple monte carlo algorithm for side chain optimization in proteins: application to model building by homology,” Proteins: Structure, Function, and Bioinformatics, vol. 14, no. 2, pp. 213–223, 1992.

[36] P. Rotkiewicz and J. Skolnick, “Fast procedure for reconstruction of full-atom protein models from reduced representations,” Journal of computational chemistry, vol. 29, no. 9, pp. 1460–1465, 2008.

[37] F. Fuchs, D. Worrall, V. Fischer, and M. Welling, “Se (3)-transformers: 3d roto-translation equivariant attention networks,” Advances in Neural Information Processing Systems, vol. 33, pp. 1970–1981, 2020.

[38] J. Kyte and R. F. Doolittle, “A simple method for displaying the hydropathic character of a protein,” Journal of Molecular Biology, vol. 157, no. 1, pp. 105–132, 1982.

[39] S. Ruder, “An overview of gradient descent optimization algorithms,” 2017.

